# Insights into the evolution of dermal armour: osteoderms in a mammal, the spiny mouse, *Acomys*

**DOI:** 10.1101/2022.10.17.512575

**Authors:** Malcolm Maden, Trey Polvadore, Arod Polanco, William B. Barbazuk, Edward Stanley

## Abstract

Osteoderms are bony plates which develop in the dermis of the skin of vertebrates, most commonly found in fishes and reptiles. They have evolved independently at least eight times in reptiles suggesting the presence of a gene regulatory network which is readily activated and inactivated. The absence of osteoderms in birds and mammals, except for the one example of armadillos, has prevented a comparative molecular approach to their evolution. However, following CT scanning, we have discovered that in two genera of *Deomyinae*, the spiny mouse *Acomys* and the brush-furred mouse, *Lophuromys* there are osteoderms present in the skin of their tails. We have studied osteoderm development within the dermis of the tail in *Acomys cahirinus* to show that they begin development before birth in the proximal part of the tail skin and they do not complete differentiation throughout the tail until 6 weeks after birth. This has allowed us to study the cellular differentiation of the osteoderms with histology and immunocytochemistry and perform RNA sequencing to identify the gene networks involved in their differentiation. There is a widespread down-regulation of keratin genes and an up-regulation of osteoblast genes and a finely balanced expression of signaling pathways as the osteoderms differentiate. Future comparisons with reptilian osteoderms may allow us to understand how these structures have evolved, why they are so rare in mammals and how they are position-specific.

## Introduction

The bones in the vertebrate skeleton develop from two independently derived pathways: the majority of the postcranial skeleton and chondocranium is endochondral, mineralizing after a cartilaginous model has been first formed, while the bones of the dermatocranium (the skull roof, jaw) and clavicles develop intramembranously by direct ossification within a matrix of collagen. The origins of these two pathways can be traced back to the earliest jawless fish. Ostracoderms and their jawed descendants, Placoderms, were covered in bony plates of dermal or integumentary bone, which represented the dominant bony structures of these organisms, and are considered to be the evolutionary precursor to the bones of the dermatocranium (1, 2). The endoskeleton of these early vertebrates was entirely cartilaginous, mineralizing in Osteichthyes and forming the endochondral skeleton. The relative proportions of these two bone-forming systems reversed early in the evolutionary history of bony vertebrates, with a reduction of dermal skeleton and the endoskeleton mineralizing and taking on a more dominant role to permit a greater degree of movement. Nevertheless the dermal skeleton has been retained in the bones of the vertebrate dermatocranium and the scales of modern day fish, which have a plate of dermal bone, much reduced in size from the ancestral form, but derived from a common ancestral integumentary element (3). Postcranial dermal bones are retained in several lineages of tetrapods in the form of osteoderms (4). These structures are common in lizards (5, 6), crocodiles (7, 8) turtles (9), and some families of anuran amphibians (10). Osteoderms are entirely absent in birds, but are known in one genus of Therapod dinosaur, *Ceratosaurus (*11), as well as from numerous Ornithischian and Sauropod dinosaurs. Dermal skeletal elements are broadly absent in Mammals, but occur in several Xenarthrans, including modern Armadillos and extinct Gyptodons and giant ground sloths(12-14). Osteoderms have been independently lost and re-evolved again with different designs at least sixteen times in amniotes (Pareiasauria, Pantestudines, Archosauria—independently evolving in pseudosuchians, thyreophorids, titanosaurs, and therapods—, Placodonta, Cingulata, and seven independent origins in Squamata—three divergent lineages of gecko, lacertids, scincoids, anguimorphs, and one species of dwarf Chameleon) so they are clearly not homologous structures (15).However their common features such as the ability of groups of cells to undergo mineralization within a well-structured dermis and having a hierarchical structure with collagen fibers joining more rigid units, thereby increasing flexibility without significantly sacrificing strength (16), may represent a deep developmental homology.

Non-avian reptiles, which have an integument covered with keratinized scales and scutes, have re-evolved osteoderms an inordinate number of times. Mammals, having evolved hairs, seem to have taken a different route and have undergone modifications of the epidermis rather than the dermis for protection. Thus we see the modified guard hairs of hedgehogs, porcupines and echidnas, the ectodermal scales of pangolins or the keratinized horns of many herbivores. Except for the armadillo and their extinct armored relatives in which osteoderms are said to be unique (13). The absence of a readily accessible and genetically tractable mammalian model of osteoderm development has seriously inhibited our understanding of their evolution and the elaboration of the gene regulatory networks involved. However, we can now make progress with an evolutionary understanding of osteoderms because we have discovered the unique phenomenon of the presence of osteoderms in the tails of two rodents in two closely related genera of *Deomyinae*, the spiny mouse, *Acomys* and the brush-furred mouse, *Lophuromys* using CT scanning. Using *Acomys cahirinus* from our in-house colony we describe the structure and development of osteoderms in the tail which develop in a proximal to distal and dorsal to ventral sequence over a 6-week period postnatally. We use histological, immunocytochemical and molecular analyses including RNA sequencing to identify which gene regulatory networks are involved in their development which together suggests a deep molecular homology exists in the mammalian dermis which can be activated at any time in evolution.

## Results

### Identification of osteoderms by CT scan

CT scans from adult museum specimens of both *Lophuromys* and *Acomys* revealed a series of dense, imbricate structures embedded in the caudal dermis (Fig. 1A-B). These structures comprise a series of 8-11 rectangular plates arranged in rings around the tail. Internal densities of the proximal structures are similar to that of the long bones, and the structures decrease in density as they approach the distal tip of the tail. No internal vascularization was revealed by the CT scans, though osteocyte lacunae were present in both proximal and distal osteoderms.

**Fig. 1.**
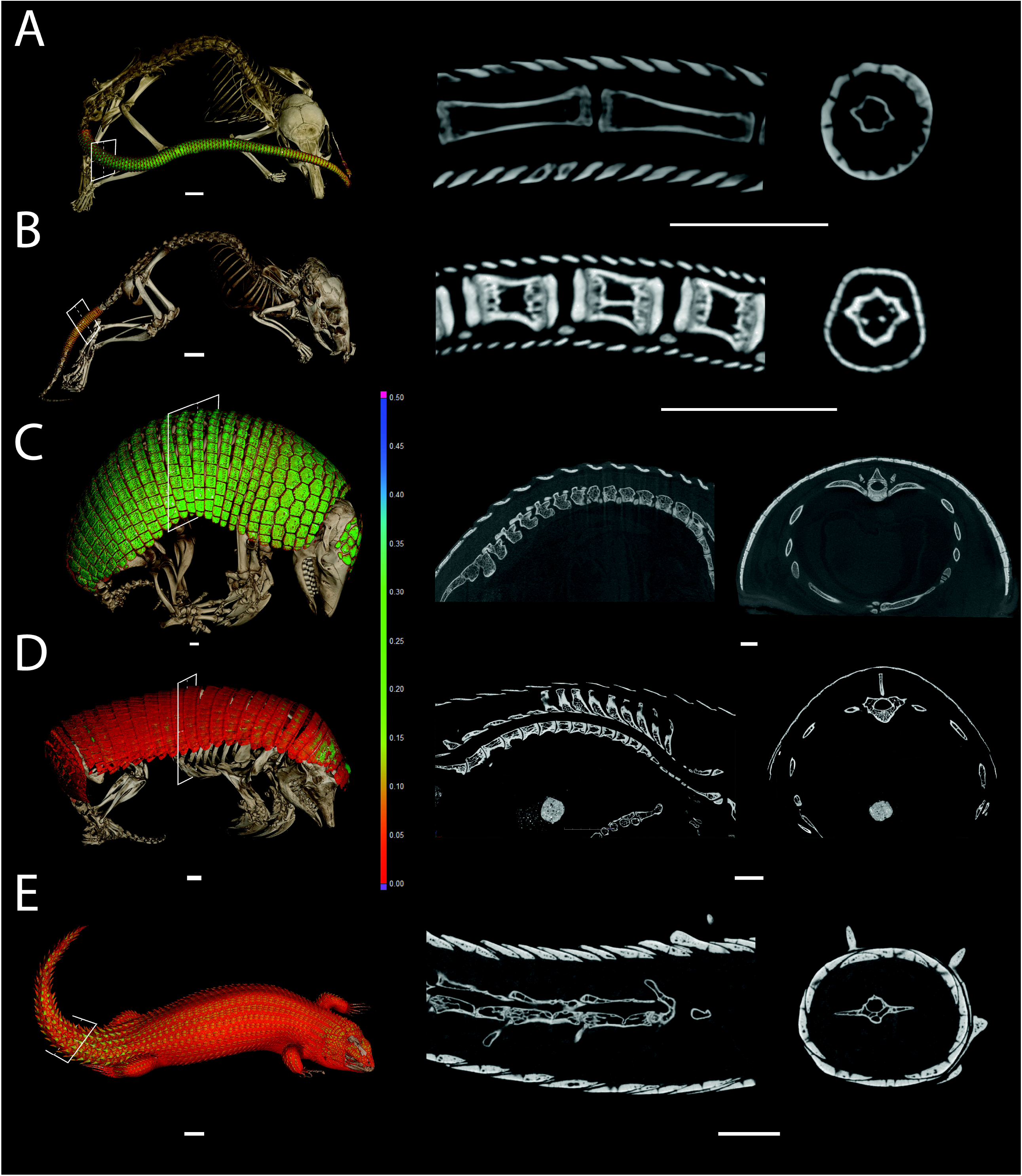
CT based, 3D renders of skeletons and osteoderms, sagittal tomograms, and transverse tomograms of **A**, *Acomys cahirinus* UF:Mammals:29706, **B**,*Lophuromys flavopunctatus* ummz:mammals:114774, **C**, *Cabassous chacoensis* UF:mammals:20650, D, Chlamyphorus truncatus fmnh:mammals:39468 and **E**, *Egernia hosmeri* UF80342. Osteoderms colorized by thickness. White squares on 3D renders denote the position of the transverse tomograms.

For comparison, scans of the armadillo (Fig. 1C) and two other squamates were obtained (Fig. D-E) which revealed a similar arrangement of dermal overlapping plates to the *Lophuromys* and *Acomys* osteoderms. In particular the tail of the squamate (Fig. 1E) looked strikingly similar to these rodent tails.

A developmental series of postnatal *Acomys* specimens beginning immediately after birth was then scanned (Fig. 2A). This revealed the presence of osteoderms in the proximal part of the tail and their absence in the distal part of the tail. Osteoderms were not present throughout the tail including the distal tip in these scan images until at least 6 weeks of age.

**Fig. 2.**
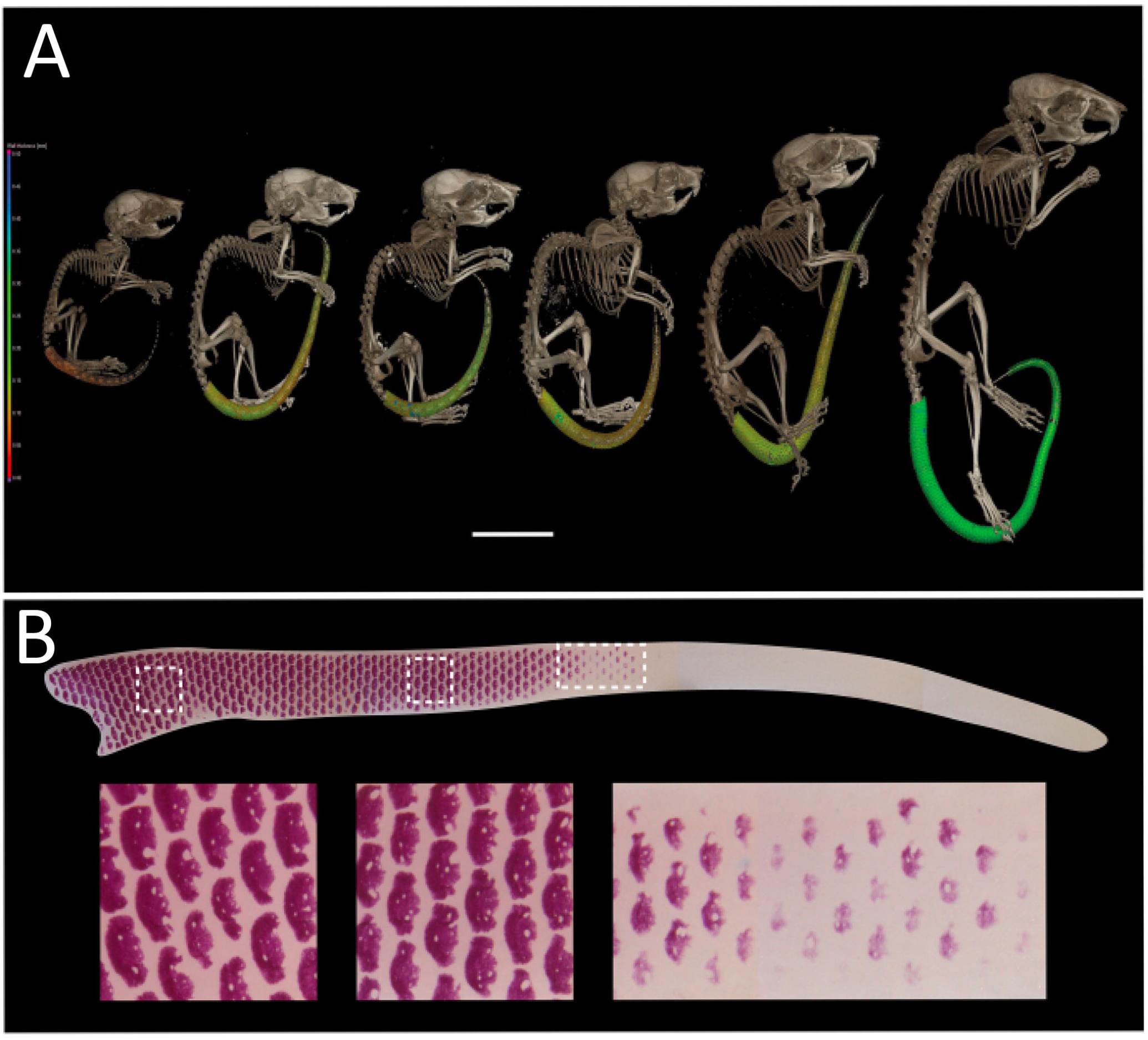
**A**, timed series of scans of *Acomys cahirinus* from post-natal day 1 (far left) to 2 year adult (far right) showing the presence of osteoderms initially only in the proximal part of the tail and then spreading throughout. **B**, Alizarin red stained newborn tail (upper) showing the presence of osteoderms in the proximal half and their absence in the distal half. A close-up of each white box is shown below revealing their regular patterning (with holes in each osteoderm) and their pattern of differentiation starting on the dorsal surface and spreading ventrally.

The absence of osteoderms in the distal newborn tail was confirmed using Alizarin red stained preparations (Fig. 2B). Here it can be seen that osteoderms appear in a gradient of differentiation beginning dorsally with one or two regions of differentiation and then their appearance progresses circumferentially to the ventral surface while the original ones on the dorsal surface expand and grow (Fig. 2B). There is therefore a gradient of differentiation both from proximal to distal and from dorsal to ventral. It was also apparent that the differentiated osteoderms contained one or more areas of non-differentiation (Fig. 2B, left panels) which presumably become the canaliculi seen in the adult osteoderm (Fig. 4H – J).

### Post-natal development of osteoderms

A developmental series of tail skin from newborn to 6 weeks of age was used to cut sections through the proximal third, the middle third and the distal third. The newborn tail had osteoderms present in the proximal third (Fig. 3A) and middle third (Fig. 3B) of the tail, but they were absent in the distal third of the tail (Fig. 3C). The same situation was present in 4 day old tails (not shown), 1 week old tails (not shown) and 2 week old tails (Fig. 3D-F). It was not until 6 weeks of age when approaching sexual maturity that osteoderms first appeared in the distal third of the tail as thin plates of bone (Fig. 3I). By this time the proximal osteoderms had grown considerably in length and particularly in thickness (Fig. 3H). For this postnatal period of time, therefore, there was a gradient of osteoderm development along the tail which allowed us the opportunity to study their histological and molecular characteristics as osteoderms developed.

**Fig. 3.**
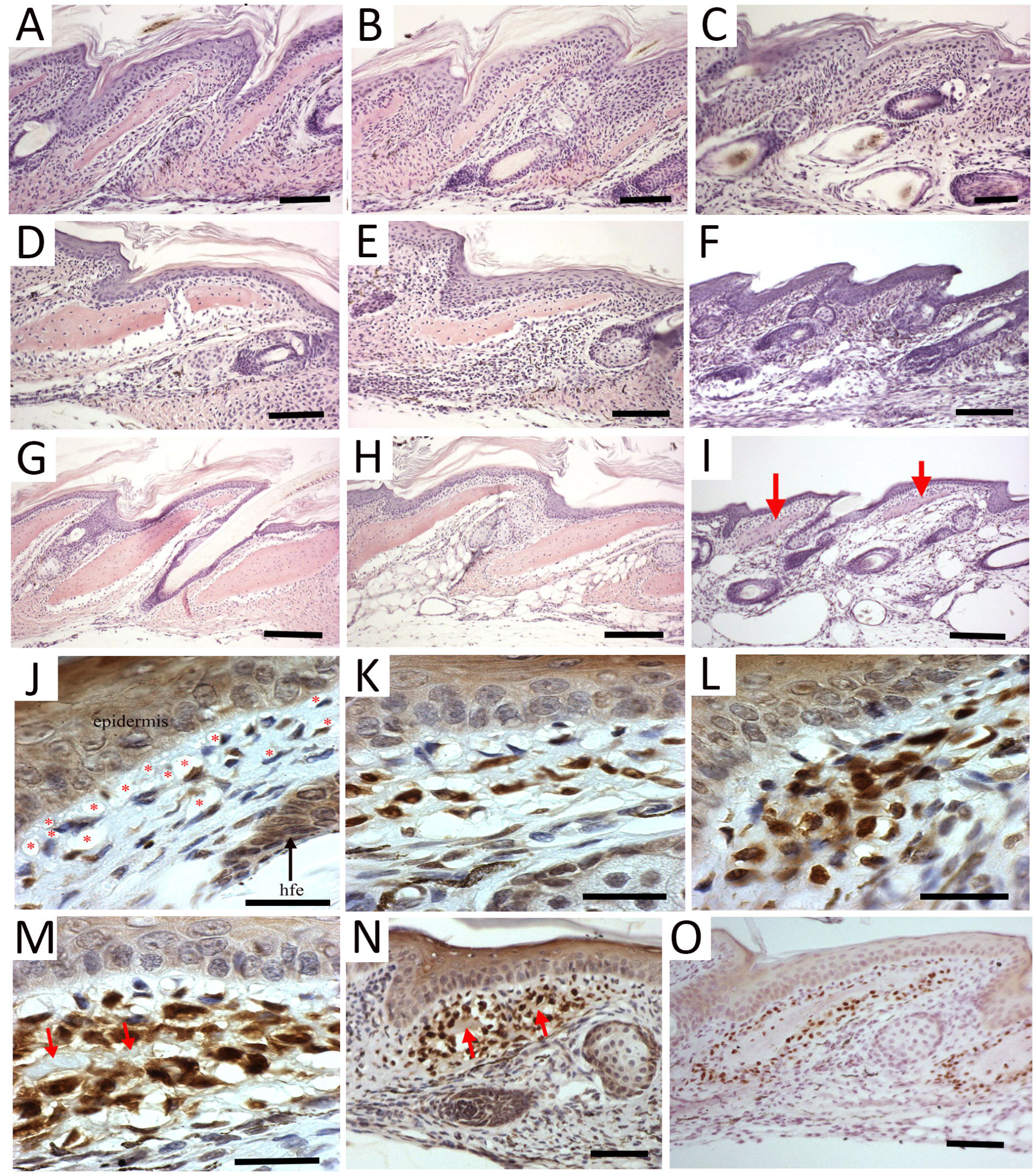
Development of *Acomys* osteoderms. **A-C**, sections through newborn tail skin showing osteoderms in the proximal (**A**), middle (**B**), but not the distal third (**C**). **D-F**, sections through 2 week-old tail skin showing osteoderms in the proximal (**D**), middle (**E**), but not the distal third (**F**). **G-I**, sections through 6 week-old tail skin showing osteoderms in the proximal (**G**), middle (**H**) and their first appearance in the distal third (**I**, red arrows). **J–O**, Osteoderm development with Ost immunocytochemistry. **J**, Ost+ve dermal cells first appear scattered in the dermis surrounded by large spaces in the dermal matrix particularly under the epidermis (red stars). Hfe = hair follicle. **K**, strong Ost+ve staining is present in cells aggregating in the middle of the dermis between the epidermis and hair follicle. Large spaces are still seen immediately below the epidermis. **L**, more Ost+ve cells appear and clump together in the dermis. **M**, a matrix (red arrows) appears in the centre of the Ost+ve cells creating a central space. **N**, lower power view showing the aggregation of Ost+ve cells with a central matrix (red arrows) being secreted. **O**, fully formed osteoderms surrounded by Ost+ve cells and Ost-ve cells in the centre surrounded by the bony matrix. A-I bar = 100μm; J-M bar = 30μm; N-O bar = 100μm.

We used an antibody to Osterix (the gene *Sp7*) which is a transcription factor involved in the development of osteoblasts, to understand how and where these osteoderms develop. The first appearance of Ost +ve cells is in individual dermal fibroblasts below the epidermis which are sandwiched between the external epidermis and the ectoderm of the hair follicle (Fig. 3J). These Ost+ve cells increase in number and intensity of staining (Fig. 3K) within a dermal matrix which seems abnormally highly vacuolated, some of which, but not all, are capillaries (red stars in Fig. 3J). These vacuoles which are seen at all stages of osteoderm development are particularly noticeable immediately underneath the epidermis (Fig. 3J - M) perhaps suggesting a role for the epidermis in the induction of the Ost +ve osteoblasts. The number of Ost +ve cells increases over time and they aggregate to form an elongated and widening accumulation of cells (Fig. 3L, M) which at its proximal region begins to secrete an extracellular matrix (red arrows in Fig 3M and N). As the matrix continues to be synthesized and the elongated and thin shape of the osteoderm appears, the osteoblasts within the matrix cease to express Osterix whereas those at the edge (presumably continuing to secrete bone matrix) continue to express Osterix and generate a ring of +ve cells around the osteoderm (Fig. 3O).

### Histological structure of adult osteoderms

Adult *Acomys* have osteoderms along the whole length of their tails and they are arranged in overlapping plates with each plate having two awl hairs (thicker ones) and a central guard hair (thinner one) emerging below it (Fig. 4A). In Alizarin red preparations the osteoderms generate rings of bone each with a slight overlap to the next ring distally to it and an offset of half an osteoderm in the circumferential axis (Fig. 4B) and the canaliculi can readily be seen (Fig. 4C). This arrangement creates a complete circle of bone (Fig. 4D) surrounding the vertebrae and spinal cord of the tail. In a section through the longitudinal axis (Fig. 4E) the regular sequential arrangement of osteoderms with hairs emerging between them can readily be seen.

**Fig. 4.**
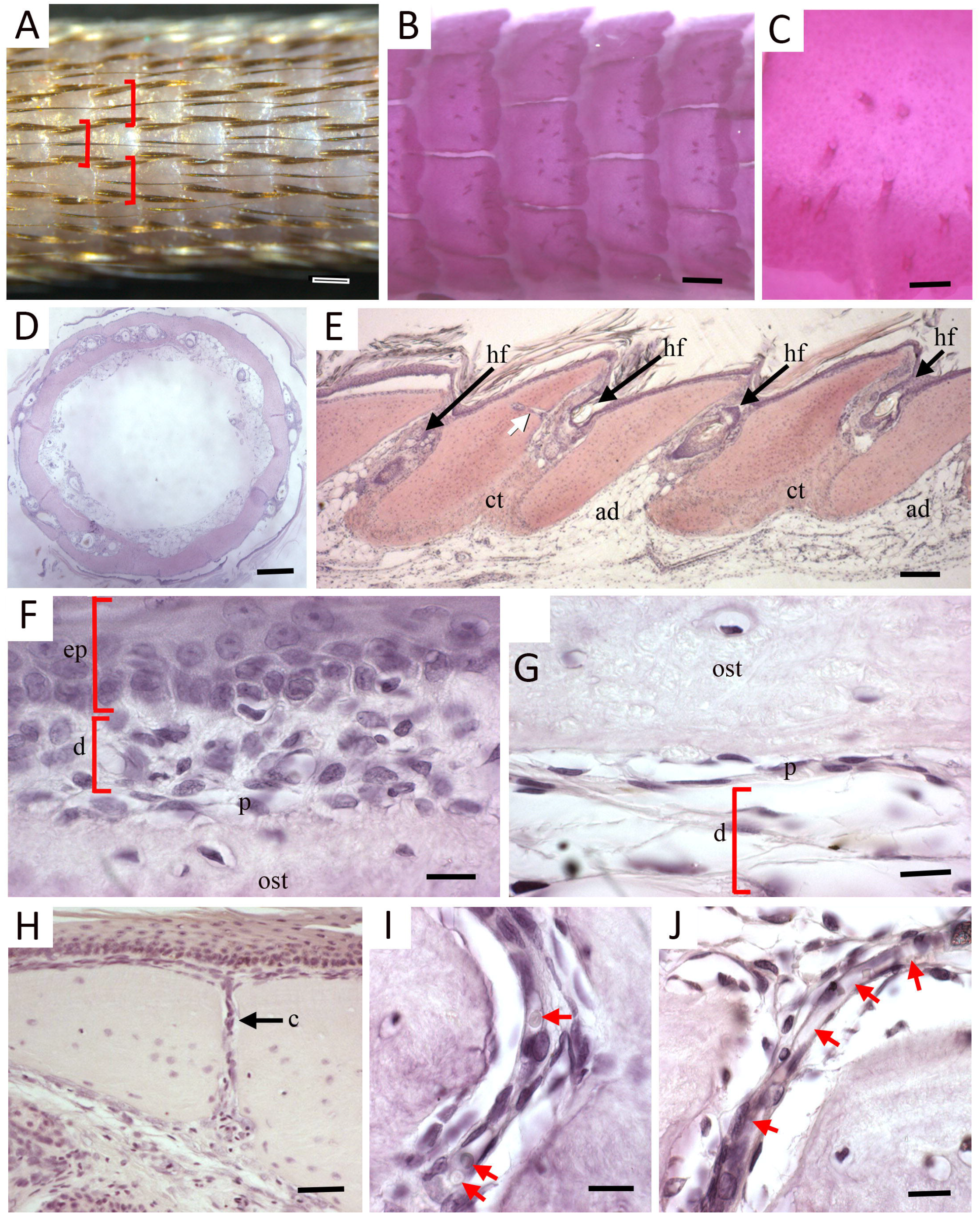
Adult *Acomys* osteoderms. **A**, external view of the adult *Acomys* tail showing the regular grouping of 3 hairs (brackets) appearing underneath an osteoderm. The 2 large hairs are awl hairs and the smaller central hair of the three is a guard hair. **B**, Alizarin stained preparation of the adult *Acomys* tail showing the formation of a continuous circle of dermal bone with each concentric ring slightly overlapping the adjacent one. Also seen are holes (canaliculi, darker dots) in each osteoderm. **C**, high power view of an Alizarin stained osteoderm showing the canaliculi. **D**, transverse section of the skin removed from the adult Acomys tail showing how the osteoderms form complete rind of bone in the dermis. **E**, longitudinal section through the skin of the adult Acomys tail showing that the osteoderms are immediately below the epidermis, the hair follicles are between each osteoderm (hf) and the osteoderms are either bound together by connective tissue (ct) of are adjacent to adipose tissue (ad) depending on the relative position of two osteoderms. The white arrow shows a canaliculus. **F**, High power view of the relation between the epidermis (ep) and the dorsal surface of the osteoderm (ost) below showing that there is a 2-3 cell layer of dermal cells (d) between and that the external cell layer of the osteoderm resembles a periosteum (p). **G**, the ventral surface of the osteoderm (ost) has far fewer dermal cells present (d) below it and the periosteum consists of flattened cells (p). **H**, section through an adult osteoderm showing a canaliculus © passing through it from dorsal to ventral. **I**, high power view of a canaliculus showing red blood cells (red arrows) in a capillary entering a canaliculus. **J**, another view of a capillary entering a canaliculus identified with red arrows. D – J, haematoxylin and eosin staining. A-B bar = 500μm; C bar = 100μm; D-E bar = 50μm; F-G bar = 20μm; H bar = 250μm; I-J bar = 20μm.

Mature osteoderms are clearly cellular rather than the acellular scales of fish (Fig. 4E, H). The dorsal surface of each osteoderm is adjacent to the epidermis of the tail with a 3-4 cell thick dermis in between (Fig. 4F) and a flattened cell layer on the osteoderm surface resembling the periosteum of more typical bones (p in Fig. 4F). The ventral surface of the osteoderm has a similarly flattened cell layer with a definite resemblance to a periosteum and below that is a loose connective tissue of the dermis (Fig. 4G). The ventral surface of the osteoderm is adjacent to either the adipose cells of the hypodermis (ad in Fig. 4E) or embedded in the connective tissue of the dermis (ct in Fig. 4E) depending on the mediolateral plane of section. It can be seen that this dermal connective tissue binds two adjacent osteoderms together (ct in Fig 4E). In between each osteoderm a hair follicle is present (hf in Fig. 4E). Canaliculi can also be seen running through the osteoderms (white arrow in Fig. 4E) which are complete channels from the dorsal to the ventral surface (c in Fig. 4H) and the canaliculi contain capillaries (Fig. 4I, J), perhaps functioning to provide the metabolic requirements of the osteoblasts within the osteoderm or to enable the vasculature to reach the layer of dermis between the dorsal periosteum and the epidermis.

### Molecular development of osteoderms

To identify the molecular players in the development of osteoderms, we performed a transcriptome analysis using bulk RNA-sequencing. Newborn *Acomys* tails were divided into 3 equal parts – proximal (with osteoderms), middle (osteoderms developing), distal (no osteoderms) and the proximal vs distal segments assayed in 6 replicates. The skin was removed from the underlying vertebrae and used for extracting total RNA having first removed a segment of each sample for histology to confirm the presence (proximal skin) or absence (distal skin) of osteoderms. Sequencing libraries were prepared from the extracted total RNA of each skin sample and sequenced on an Illumina NovaSeq6000 platform, yielding approximately 898 million paired-end, 150 bp length reads. Analysis using fastQC indicated that all 12 libraries passed standard quality control assessments following trimming of low-quality reads and adapter sequences, and all libraries were utilized in downstream analyses.

A transcriptome cataloging gene expression in the developing spiny mouse tail was assembled using the *Acomys* skin transcriptome (accession #GSE113081 (17)) as a reference. HISAT2 (18) and StringTie (19) were used to align and assemble the trimmed paired-end reads, respectively, and featurecounts used to determine counts per transcript (Supplemental Table S1). Counts for transcripts sharing the same Trinity cluster and gene designation were collapsed essentially converting counts/isoform to counts/gene (Supplemental Table S2). Differential expression analysis using edgeR with a paired sample design in the gene level counts data identified 4366 genes with differential expression in proximal and distal tail skin, with 1814 genes downregulated and 2552 upregulated at an adjusted p-value of 0.05 (Supplemental Table S3, Supplemental Figure S2). The counts data converted to log2 CPM (counts per million) for genes exhibiting significant differential expression (pvalue >= 0.05) and log2FC >= 2 were hierarchically clustered and exhibited as a gplots heatmap2 plot (Supplemental Figure S3). Hierarchical clustering visualizing differences in gene expression between proximal and distal skin across all significantly differentially expressed genes (Supplemental Figure S1) separates treatment groups (proximal/distal) but indicates variation between individuals.

The top 20 up-regulated (‘+’ log2FC) and the top twenty down-regulated (‘-’ log2FC) genes between *Acomys* proximal vs. distal tail skin samples that can be annotated by similarity to Mus orthologues are shown in Table 1. These counts data converted to log2 CPM (counts per million) were hierarchically clustered and exhibited as a gplots heatmap.2 function (Fig. 5). Inter-sample variation in gene expression is clear in both the heatmap of all significant genes exhibiting a lof2 fold change of <=2 and the heatmap displaying hierarchical clustering of the top 20 up and top 20 down regulated transcripts. A non-clustered heatmap (supplemental Figure S4) of the top 20 up and 20 down regulated genes data displaying paired proximal – distal samples side by side and gene order from top up to top down (same order as Table 1) further illustrates the intersample variation but makes clear that the trends in genes expression between distal and proximal pairs are consistent between samples.

**Table 1.** Top 20 up-regulated and the top twenty down-regulated genes between Acomys Proximal vs. Distal tail skin samples annotated by Mus orthologues.

| DOWN-REGULATED |  |  | UP-REGULATED |  |  |
| --- | --- | --- | --- | --- | --- |
| transcript | gene name | Log2FC | transcript | gene name | Log2FC |
| TR50180\|c0_g3 | *Krt71* | -6.08 | TR91285\|c0_g1 | *Bglap3* | 5.21 |
| TR152503\|c0_g3 | *Krt28* | -6.05 | TR53845\|c0_g1 | *Ibsp* | 4.75 |
| TR152503\|c0_g2 | *Krt27* | -6.02 | TR91530\|c0_g1 | *Fat3* | 4.56 |
| TR36557\|c0_g1 | *Krt73* | -5.68 | TR108990\|c0_g1 | *Phex* | 4.42 |
| TR68726\|c0_g2 | *Krt31/Krt33b* | -5.65 | TR160427\|c0_g1 | *Spp1* | 4.35 |
| TR152503\|c0_g4 | *Krt25* | -5.52 | TR1455\|c0_g1 | *Fam43b* | 4.23 |
| TR121791\|c0_g1 | *Fabp9* | -5.35 | TR45613\|c0_g1 | *Panx3* | 4.23 |
| TR151232\|c0_g2 | *Gprc5d* | -5.28 | TR3405\|c0_g1 | *Ostn* | 4.14 |
| TR150451\|c0_g1 | *Krt72* | -5.18 | TR157631\|c0_g1 | *Dkk1* | 4.13 |
| TR47183\|c0_g2 | *Tchh* | -5.11 | TR122953\|c0_g1 | *Dlg2* | 4.01 |
| TR78826\|c0_g1 | *Krtap19-1* | -5.02 | TR56704\|c0_g2 | *Sp7* | 3.85 |
| TR14881\|c0_g2 | *Krt34* | -5.00 | TR141673\|c0_g1 | *Slc13a5* | 3.84 |
| TR138942\|c0_g1 | *Tnnc1* | -4.90 | TR57247\|c0_g1 | *Ifitm5* | 3.83 |
| TR76618\|c2_g1 | *Fam26d* | -4.75 | TR36614\|c0_g1 | *Omd* | 3.78 |
| TR42445\|c0_g1 | *Krtap1-4* | -4.74 | TR106888\|c0_g1 | *Slc8a3* | 3.73 |
| TR76306\|c0_g1 | *Krtap14* | -4.63 | TR87506\|c0_g1 | *Slitrk1* | 3.70 |
| TR38697\|c0_g1 | *Krtap5-5* | -4.62 | TR30280\|c0_g1 | *Col22a1* | 3.63 |
| TR91685\|c0_g1 | *Krtap15* | -4.39 | TR21906\|c1_g1 | *Tmppe* | 3.61 |
| TR68726\|c0_g1 | *Krt33a/Krt34* | -4.26 | TR29945\|c3_g1 | *Lep* | 3.58 |
| TR109908\|c0_g1 | *Tnni1* | -4.19 | TR1115\|c0_g2 | *Cthrc1* | 3.51 |

**Fig. 5.**
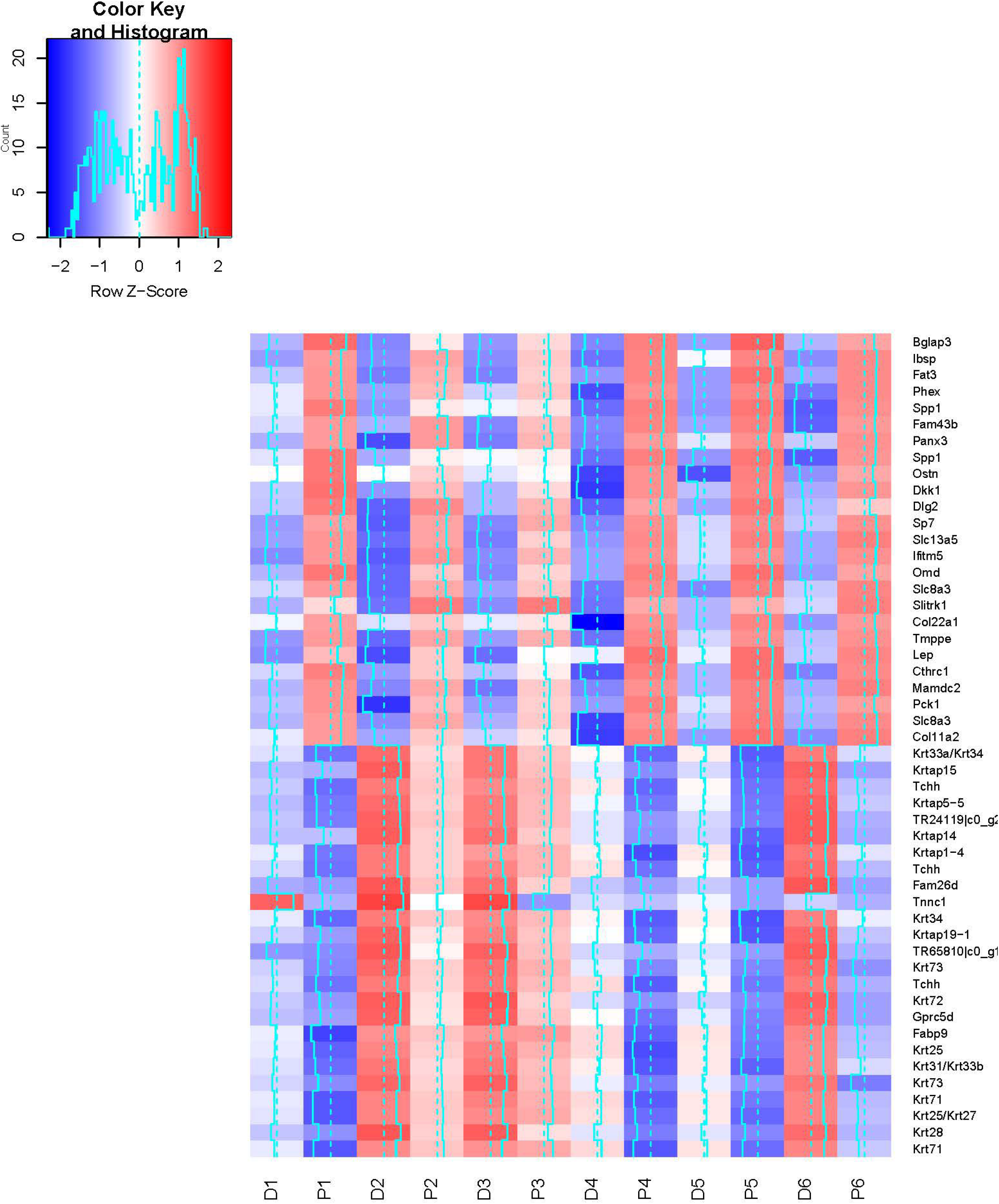
Heatmap of the top 25 (largest log2 FC) variable transcripts between proximal and distal newborn *Acomys* tail skin. The six distal skin samples are D1 – D6 and the six proximal samples are P1 – P6 and these samples separate well.

What is strikingly apparent in the proximal tail skin is the switch from hair formation to bone formation. Of the top 20 down-regulated genes with annotations in the proximal tail skin, 16 are associated with hairs. 7 are hair keratins (*Krt71, Krt 28, Krt25, Krt27, Krt73, Krt72, Krt34*), 2 are cuticular keratins (*Krt33b, Krt 33a*), 5 are Keratin associated proteins (*KRTAP19-1, 1-4, 14, 5-5, 15*) and one more gives strength to hairs (*Tchh*), while FabP9 is expressed in the internal root sheath of hair follicles. Only four genes have no apparent relationship to hairs: a G protein-coupled receptor (*Gprc5d*), Calcium Homeostasis Modulator Family Member 4 (*Fam26d)*, and two muscle genes (*Tnni1, Tnnc1)*. Interestingly the KTRAP multigene family plays an important role in hair formation and morphology and has been associated with phenotypic differences in hairs (20). In particular the spines of hedgehog which are similar to the spines of spiny mice are correlated to the number of KTRAP genes and the number of pseudogenes. It would be interesting to undertake a genomic study of these genes in *Acomys*.

The switch to bone formation is clear as 12 of the top 20 up-regulated genes are directly involved in or associated with bone as structural proteins (Bglap2, *Ibsp, Col22a1*), mineralization (Phex, *Fat, Ifitm5, Omd*) or attachment proteins (*Spp1*), hormones or transcription factors (*Ostn, Sp7, Lep*). Images of Sp7 immunocytochemistry (Osterix) can be seen in Fig. 3 confirming its up-regulation in the osteoderms of the proximal tail skin and octeocrin (*Ostn*) is a direct gene target of Sp7 in the differentiation of osteocytes (21). The other top genes are involved in Wnt signaling (*Dkk1, Cthrc1*), metabolism (*Slc13a5, Slc8a3*,), gap junctions (*Panx3*) or no known function (*Fam43b*).

Pathway analysis using IPA predicted the activation of osteoblast differentiation (p-value 7.24E-18, z-score 2.14),connective tissue differentiation (p-value 1.34E-20, z-score 2.00) and mineralization of bone (p-value 4.45E-16, z-score 2.257) in the proximal tail skin samples. All three of these developmental processes are important for the formation of bone, and the genes involved may shed light on the mechanisms that initiate and drive osteoderm development. Twenty-two genes including *Panx3* (upregulated 4.23-fold), *Sp7* (upregulated 3.85-fold), and *Runx2* (upregulated 2.31-fold) were identified by IPA as having differential expression consistent with the activation of osteoblast differentiation (Table 2). Likewise, *Areg* and *Mstn* are expected to inhibit osteoblast differentiation and their down-regulation (log2FC=-1.688 and log2FC=-1.72, respectively) in proximal vs. distal tail skin are consistent with activation of osteoblast differentiation. Indeed, *Runx2* and *Osterix* (*Sp7*) are the two essential transcription factors for osteoblast differentiation.

**Table 2.**
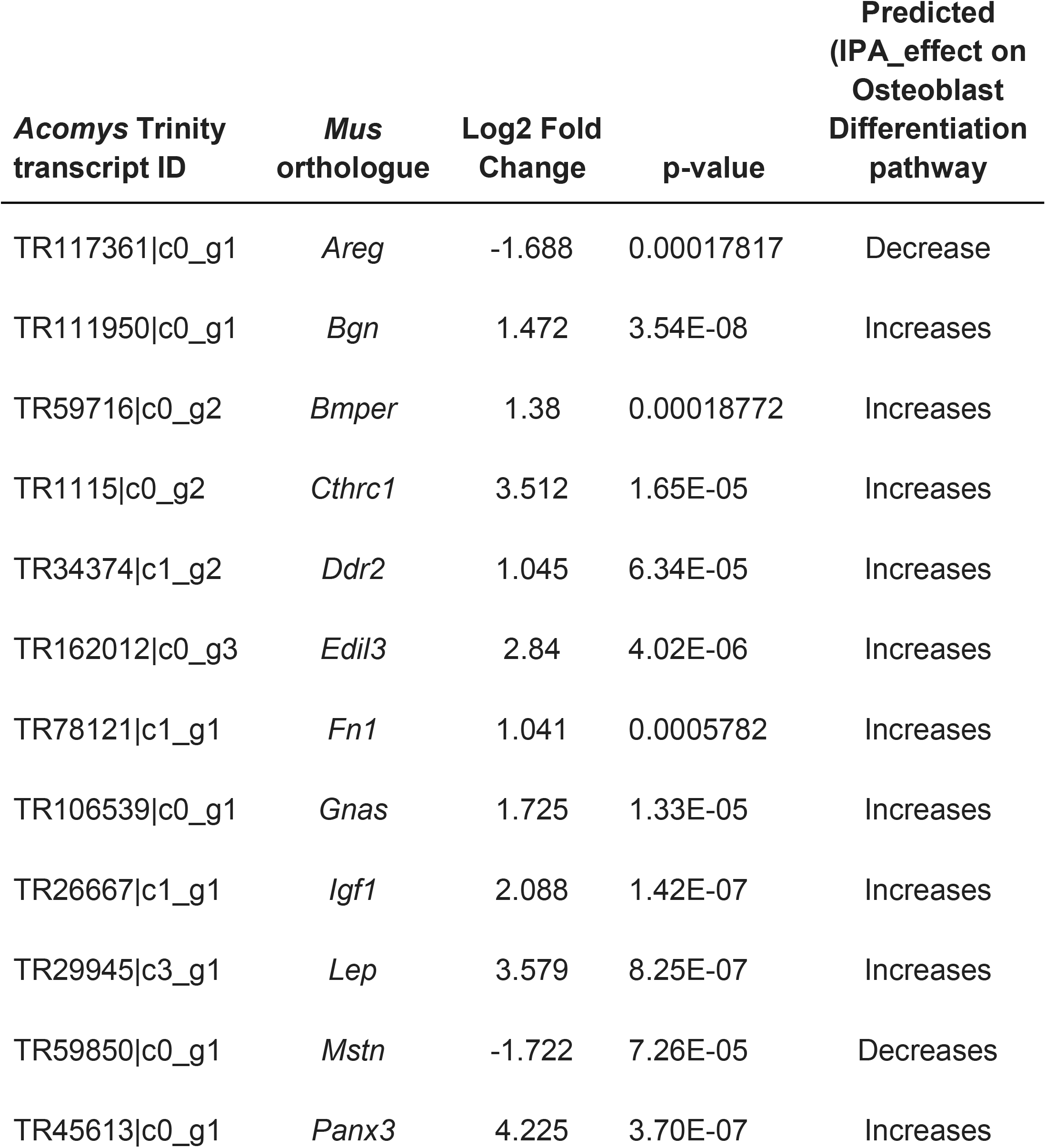
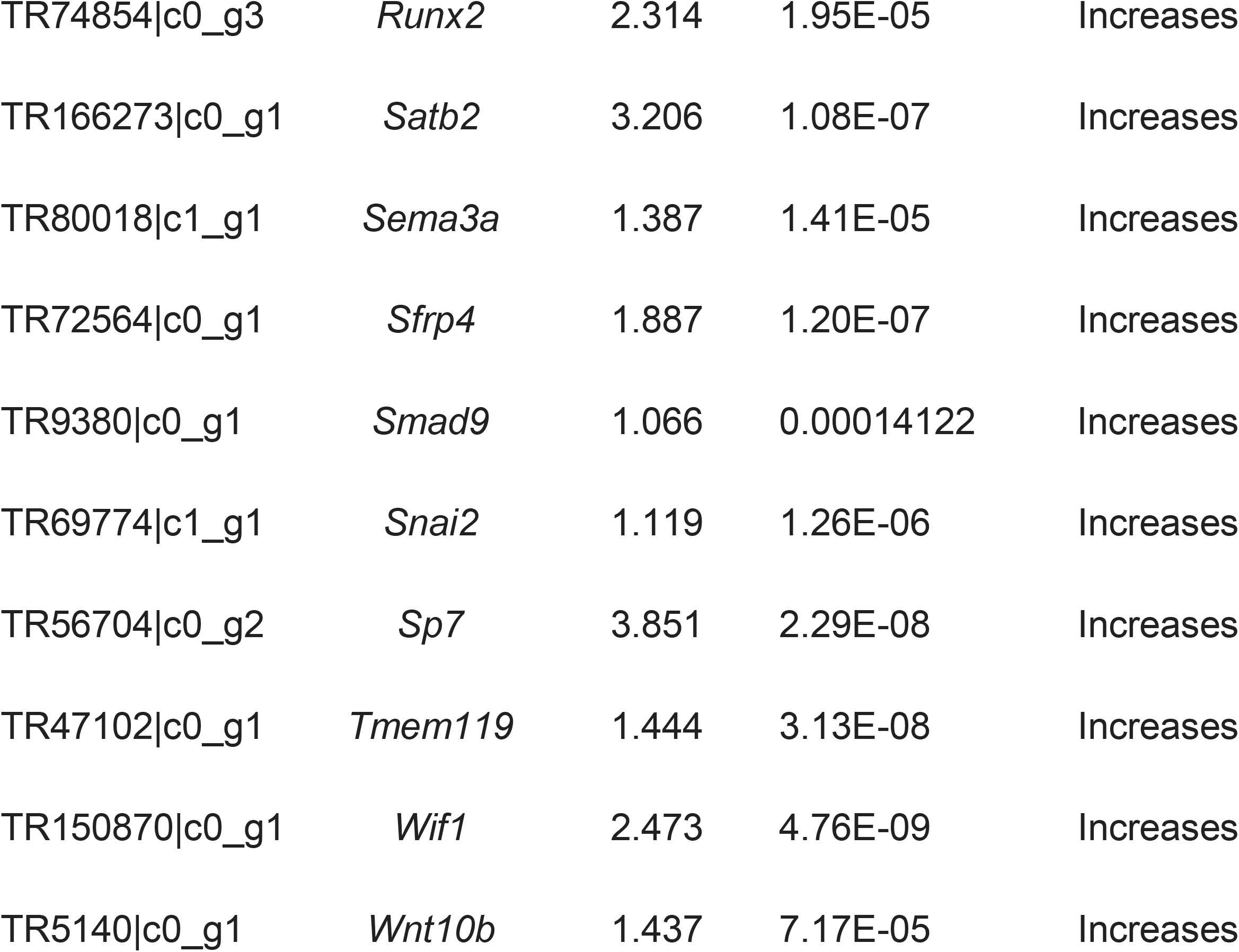
Genes with expression indicating activation of osteoblast differentiation. Column four indicates the effect expression of the gene is predicted to have on the osteoblast differentiation pathway as predicted by IPA. *Panx3* is expected to have a positive effect on osteoblast differentiation, and its upregulation (log2FC=4.225) in proximal vs. distal tail skin is consistent with activation of osteoblast differentiation. Likewise, *Areg* expression is expected to inhibit osteoblast differentiation and its down-regulation (log2FC=-1.688) in proximal vs. distal tail skin is consistent with activation of osteoblast differentiation.

We also identified 8 collagens that are upregulated at least twofold in the proximal tail (Table 3), including collagens of types 1, 3, 5, 11, 22 and 24. Additional molecular evidence for the formation of osteoderms in the proximal tail can be seen by the expression of 12 out of 33 genes cataloged by IPA as being involved in the activation of bone mineralization (p-value 4.45E-16, z-score 2.26).

**Table 3.** Collagen genes upregulated at least twofold in proximal tail skin.

| Transcript | Gene | Log2FC | P-value |
| --- | --- | --- | --- |
| TR45475\|c2_g1 | *Col1a1* | 2.79 | 1.41E-06 |
| TR160543\|c1_g1 | *Col1a2* | 2.35 | 1.07E-08 |
| TR1240\|c0_g1 | *Col3a1* | 2.28 | 7.93E-07 |
| TR7472\|c1_g2 | *Col5a2* | 2.63 | 4.59E-05 |
| TR74197\|c1_g2 | *Col11a1* | 2.1 | 2.10E-05 |
| TR468\|c0_g1 | *Col11a2* | 3.36 | 1.14E-04 |
| TR30280\|c0_g1 | *Col22a1* | 3.63 | 4.39E-04 |
| TR15949\|c0_g3 | *Col24a1* | 2.01 | 1.94E-07 |

There are five canonical pathways involved in bone formation, namely BMP, FGF, HH, Wnt and EDA pathways and we could identify four of them in this data. In the proximal gene lists we identified several BMPs, namely *Bmp1* (log2FC = 1.02), *Bmp3* (log2FC = 1.29), *Bmp6* (log2FC = 0.52) and the inhibitor *Bmper* (log2FC =1 .38). Several FGF genes were identified including, *Fgf12* (log2FC = 0.77), *Fgf13* (log2FC = 1.02), *Fgf22* (log2FC = −1.00), *Fgfbp3* (log2FC = −0.58) and *Fgf5* (log2FC = −2.97). The hedgehog pathway members *Smo* (log2FC 0.61) and *Hhip* (log2FC = 1.53) were identified along with the transcription factors *Gli1* (log2FC 1.43) and *Gli3* (log2FC = 0.60).

The Wnt pathway components were well represented and ‘Wnt-b catenin signalling’ was identified by IPA. Interestingly, both positive components and inhibitors were identified. The Wnt ligands *Wnt2* (log2FC = 1.3<u>7</u>), *Wnt10b* (log2FC = 1.44) and *Wnt6* (log2FC = 0.87) were up-regulated whereas *Wnt5a* (log2FC = 0.50) and *Wnt9a* (log2FC = −0.55 were down-regulated. The Wnt inhibitor *Dkk1* (log2FC = 4.13 and *Wif1* (log2FC = 2.47) were strongly up-regulated suggesting osteoderm development requires Wnt inhibition. Other Dkks were *Dkk2* (log2FC = 0.82) and *Dkk3* (log2FC = 1.13). The receptor components *Fzd2* (log2FC = −0.69), *Fzd3* (log2FC = 0.38), *Fzd8* (log2FC = 0.83) and *Lrp5* (log2FC = 0.54) were identified along with the secreted frizzleds *Sfrp1* (logFC = −0.33) and *Sfrp4* (log2FC = 1.89). Another Wnt pathway inhibitor *Sclerostin* was highly up-regulated 3.18 fold (log2) and *Cdh2* which interacts with b-catenin was up-regulated 2.55-fold (log2). In contrast *Wisp1* which promotes osteoblast differentiation by binding BMP2 and enhances BMP function in osteogenesis was up-regulated 1.82-fold (log2). This data suggests that the careful *regulation* of Wnt signaling is important in the formation of osteocytes so that there is no excessive production of bone throughout the dermis. This is discussed further below.

## Discussion

We reveal here a unique phenomenon in rodents – the presence of osteoderms in the tail of 2 species of *Deomynidae*, the spiny mouse *Acomys* and its close relative *Lophuryomus*. An earlier report of osteoderms in another species of *Acomys* suggests that they are present throughout these two Genera (22). It is almost a unique phenomenon in the entire class of Mammalia as the only other mammal with dermal bones is the armadillo. Since these two groups are separated by at least 70 million years it suggests that these structures have independently evolved rather than being homologous. Apart from these two unrelated mammals there are no reports of osteoderms in birds whereas in reptiles they are widespread from lizards to turtles and crocodiles and have been independently evolved at least 16 times (see Introduction). Clearly the gene regulatory network to generate dermal bones has been preserved throughout evolution and can be reactivated with the appropriate cues and it would be interesting to determine whether the other members of the subfamily, namely *Uranomys* and *Deomys* also have osteoderms in the tail.

Thanks to the presence of a breeding colony of *Acomys* which we use for regeneration studies (23-25) we could readily examine the development of osteoderms in the *Acomys* tail because at birth only the proximal third has differentiated osteoderms. Differentiation of the newly forming osteoderms begins on the dorsal surface of the tail skin and spreads distally and ventrally. In the middle third of the tail they are just beginning to form by the aggregation of Osterix +ve osteoblasts in the dermis and they are absent in the distal third of the newborn tail.

The process of osteoblast differentiation begins in a small number of dermal fibroblasts just below the epidermis and at the same time large cavities appear within the dermal matrix perhaps suggesting an epidermally secreted factor may play a role in the induction of osteoblast differentiation. The aggregation of osteoblasts then begin to secrete bone matrix themselves and assemble into a flat plate surrounded by osteoblasts and osteocytes within the matrix. The fully differentiated osteoderm has canaliculi passing through it, perhaps to allow the passage of blood vessels as these can be seen passing through the larger gaps. At their internal surface the osteoderms are attached to thick connective tissue joining adjacent ones and creating a strong ring of imbricate scales and each ring is offset by half an osteoderm relative to the adjacent one showing a very precise patterning event has taken place during development. Furthermore, they are precisely arranged relative to the hairs of the tail so that 3 hairs emerge below each osteoderm. If induction does occur due to an epidermal signal it would be fascinating to examine the patterning relationship between the hairs (epidermal placodes) and osteoderm inducing signal much earlier in tail development.

RNA seq revealed the up-regulation of a large number of genes involved in osteoblast differentiation, connective tissue differentiation and activation of bone cell differentiation, which comprised the vast majority of the top 20 up-regulated gene in proximal tail skin. Conversely there was a corresponding down-regulation of a large number of keratin genes. The up-regulated genes included many collagens, mineralization genes, structural protein genes, transcription factors and secreted factors. The *Runx2* - *Sp7* – *Osteocrin* induction sequence was operative here and *Osteocrin* is a secreted protein which promotes osteocyte dendrite formation and maintenance (21). Some less well-known factors involved in bone formation that are upregulated in proximal vs. distal tail skin samples include leptin (log2FC=3.56; Supplemental Table 4) and the leptin receptor (log2FC=2.41; Supplemental Table 4), which are both involved in osteoblast proliferation and differentiation (26). The induction of some of these factors in dermal fibroblasts may play a role in osteoderm generation and now we have a mammalian system to work with we can test the efficacy of these individual genes we have identified by transfecting dermal fibroblasts. What these future experiments will not tell us, however, is the fascinating ‘positional’ question of why osteoderms only appeared in the tail and not all over the body.

Genes of the Wnt pathway which are also involved in bone development (27) were also up-regulated, but surprisingly so are genes which are antagonistic to the Wnt pathway such as *Dkk1, Wnt inhibitory factor, Sfrp4* and *Sclerostin*. It is reasonable to assume therefore that careful control of the Wnt pathway by the balance between inducers and inhibitors are important for the precise differentiation of osteoderms so that the correct size of the osteoderm primordium is generated and they do not overgrow as they are precisely arranged in the circumferential and longitudinal axis. Indeed, mutations in each of these Wnt inhibitors identified here (28-32), results in increased bone density and mass demonstrating their role in the control of bone growth.

Despite considerable structural variation across the vertebrates, osteoderms have two common features: their origin is within the dermis and structurally they are composed of osseous tissue without the formation of a cartilaginous precursor as is the case in endochondral ossification. As such it is suggested that osteoderms share a common evolutionary origin (3, 4) stretching back to the Placoderms. Thus the origin of the cells that give rise to the wide array of dermal armor and osteoderms across the vertebrates is of considerable evolutionary interest and has been hypothesized to be of neural crest origin because of the similarity to tooth formation and because the development of osteoderms by direct ossification of mesenchyme resembles intramembranous ossification of neural crest cells which takes place during the development of the skull. Although trunk neural crest does not normally make bone it has skeletogenic potential (33) and these ideas place the neural crest as a driving force in dermal armor evolution assuming the loss of their odontogenic capacity (4).

On the other hand, the posterior skull bones of both mouse and chick which also develop by intramembranous ossification are mesodermally derived (34-36) not neural crest derived, so direct ossification of mesodermal cells themselves is a normal process during development. In addition, there is no evidence for a neural crest origin of scales in zebrafish (37) providing phylogenetic evidence for the mesodermal origin of dermal bony structures. If the mouse possessed osteoderms this question could readily be answered using the neural crest lineage labelled transgenic lines which have been so instrumental in realizing the origin of cranial structures (35, 38). There seem to be no studies on neural crest in the developing mouse tail using these transgenics but since tails develop hairs there must be some neural crest cells present in the tail dermis to produce the dermal hair placodes and thus, perhaps, also generate the osteoderms. Identifying the origin of osteoderms is an extremely interesting evolutionary question which we can begin to address now that we have identified their presence in an experimentally tractable mammalian model system.

## Materials and Methods

### Animals

Animals were obtained from our colony of *Acomys cahirinus* housed at the University of Florida and used in accordance with protocols approved by the Institutional Animal Care and Use Committee (IACUC) (protocol number 202107707). Museum specimens were obtained from the Florida Museum of Natural History, Peabody Museum and the University of Michigan Museum of Zoology.

### CT Scanning

A Yale Peabody specimen of *Acomys cahirinus* (YPM-MAM-005794) was imaged as part of the Open Vertebrate (oVert) Thematic Collections Network initiative (NSF1701714). Following discovery of the structures resembling osteoderms in the caudal region of the specimen, an ontogenetic series of *A. cahirinus* was imaged. All specimens were scanned using a dual tube GE Phoenix V|tome|X M CT system at the University of Florida’s Nanoscale Research Facility (NRF). We employed the 240kV microCT tube focus tube, and modified the voltage, current, detector capture time and rotation angles to optimize contrast and signal and minimize artifacts. Radiographs were converted into tomograms using Phoenix DatosOS|X reconstruction software. Submicron voxel resolution CT scans of second series of osteoderms were imaged using an Xradia Versa 620 system at the NRF and processed in CT datasets for comparative squamate and mammalian material, including a closely related genus of deomyine mouse, *Lophuromys flavopunctatus (UMMZ-MAM-114774)*, were downloaded from Morphosource.org. All CT datasets were post-processed in VGStudio Max 2022.1 (VolumeGraphics, Heidelberg Germany) to digitally isolate regions of interest and recover volumetric measurements. Wall thickness analyses were performed on isolated osteoderm regions. All CT datasets (Tomogram stacks and metadata) are freely available to download at morphosource.org.

### Histology

Tails of various ages were fixed overnight in 4% PFA and then decalcified for at least 1 week in 14% EDTA with daily changes of solution. Tissues were processed for paraffin wax histology and sectioned at 10mm. Sections for haematoxylin and eosin staining were dewaxed to deionized water and then placed in Harris’ haematoxylin for 3 minutes and washed in running tap water for 10 minutes. Slides were then placed in 0.1% eosin in 70% alcohol for 3 minutes followed by 90% ethanol, 100% ethanol twice each for 1 minute and then twice in xylene each for 5 minutes before being coverslipped with Cytoseal.

For immunocytochemistry slides were dewaxed and rehydrated to deionized water and placed in 2% hydrogen peroxide for 30 minutes. After 5 minutes washing in deionized water antigen retrieval was performed by heating in a microwave for 4 minutes in sodium citrate buffer at pH6. Slides were then placed in PBS for 5 minutes and blocked in diluted normal serum in PBS for 1 hr at room temperature following the Vectastain Elite ABC kit procedurE. A Sp7/Osterix antibody (Abcam ab227820) was diluted at 1 in 500 in blocking buffer and applied to the slides overnight at 4°C. After washing the slides 2x in PBS the relevant biotinylated secondary antibody was diluted in blocking buffer and applied for 1 hr at room temperature followed by ABC reagent (1 hr) and DAB (2 mins) according to the Vectastain protocol. After washing the slides were lightly stained with haematoxylin (1 min), dehydrated and mounted in Cytoseal.

Alizarin red staining for whole-mount tail osteoderms was performed by fixing the tails in 10% neutral buffered formalin overnight, washing in PBS and then another overnight step in 30% hydrogen peroxide. After another wash in PBS the tails were stained for 45 minutes in Alizarin prepared by dissolving 1g of Alizarin Red S in 24 ml distilled water and adjusting the pH to 4.2 with conc HCl. They were then washed in distilled water and left overnight in 1% KOH. The specimens were then cleared in 50% glycerol in 1% KOH for 24 hrs or longer as required.

### RNA extraction

Full-thickness skin was collected from the proximal and distal tail of six newborn *Acomys* pups and either stored at −20 oC in RNALater or immediately processed for RNA extraction. Samples of 13-34 mg skin were homogenized with a handheld mechanical homogenizer, and total RNA was extracted using a RNeasy Fibrous Tissue Mini Kit (QIAGEN 74704) according to the manufacturer’s protocol. A total of 17.7-42.8 mg of RNA was extracted from each sample (means of 29.8 mg (*s* = 10.1 mg) and 22.8 mg (*s* = 2.6mg) for proximal and distal tail samples, respectively), as quantified using a NanoDrop ND-1000 spectrophotometer. Agilent TapeStation analysis assigned RINs ranging from 7.7 to 9.3 (mean RIN of 8.6, *s* = 0.63).

### Library construction and RNA-sequencing

An Illumina TruSeq stranded library was constructed via poly-A selection for each individual skin sample for a total of 12 libraries (six proximal and six distal). The libraries were sequenced on the Illumina NovaSeq6000 platform to yield nearly 898 million paired-end, 150 bp reads (mean of 74.8 M read pairs (*s* = 9.1 M read pairs) per library). Library construction and sequencing were performed by HudsonAlpha Discovery in Huntsville, Alabama, USA.

### Sequencing data processing

RNA-seq FASTQ-files were quality and adapter trimmed using Trimmomatic v0.39 (39) with the parameters : LEADING:3 TRAILING:3 SLIDINGWINDOW:4:15 MINLEN:36 and ILLUMINACLIP with the TruSeq3-PE-2.fa adapter set (available on the Trimmomatic github page: https://github.com/timflutre/trimmomatic/tree/master/adapters) and values of 2, 30,10 for seed mismatches, palindrome clip threshold and simple clip threshold, respectively. The preprocessed reads were aligned to the *Acomys* skin transcriptome (Geo Accession GSE113081) with HiSat2 (v2.2.1) (18) using default parameters for paired-end reads. HiSat output SAM files were converted to BAM format and sorted with SamTools (1.12) (40) and used as input to Stringtie (19). Individual read set alignments representing the output from HiSat2 were assembled into transcripts with Stringtie using default settings. Output GFF files were merged with Stringtie called a second time with the following parameters: --merge. Stringtie was called a third time using the individual sorted BAM file outputs of HiSat and the following parameters: -G and the merged GFF as reference, and -A to report transcript abundance.

### Assessment of gene expression and identification of gene differentially expressed between distal and proximal *Acomys* tail skin samples

The *Acomys* transcriptome (Geo Accession GSE113081) is a Trinity (41) assembly described in Brant et al. 2019 (17). Trinity assembles in a hierarchical manner first assembling RNA-Seq reads into contigs and grouping contigs that have sequence similarity. The contig groups are used to construct De Brujin graphs that are traversed to identify transcript isoforms. The names of the individual transcript assemblies outputted by Trinity are structured to capture the cluster ID (related contigs) the transcript was constructed from, the group (gene) within the cluster the transcript belongs to and a final isoform designation. Thus, each individual transcripts within the Trinity assembled Acomys transcriptome (GSE113081) can be assigned to an independent loci based on its sequence ID. Counts per transcript for each RNA-Seq sample were determined with the featureCounts function from the Rsubreads package (v2.0.0) (42) (Supplemental Table S1), which map the aligned reads described in sorted BAM HiSat2 output to individual transcripts within the *Acomys* trancriptome assembly. Featurecounts (43) was called with the following parameters: -s 2 -p -t exon -g gene_id - T 8. Counts to isoforms that are representatives of the same trinity loci were aggregated (Supplementary table 2) to enable gene level expression analysis. Brant et al. annotated the *Acomys* transcriptome and defined *Mus* – *Acomys* orthologous for a subset of the transcripts. These annotations were used in this analysis. Differentially expressed genes were discovered with the EdgeR package (v3.36) (44) using the aggregated count matrix as input. Count data were TMM normalized, distal/proximal tail skin samples from the same individual were treated as paired samples, and genes differentially expressed between *Acomys* distal and proximal tail skin samples with pvalues <= 0.05 after Benjamini & Hochberg false discovery rate correction were reported as significant (Supplemental Table 3. Log2 cpm (counts per million) values were calculated for each gene across all samples by the CPM method within EdgeR (Supplemental Table S4) and used for PCA assessment and heatmap construction. Note that the trinity transcript assembly used as a reference (Geo Accession GSE113081) was constructed without a genome reference. Thus, it is expected that transcripts represented by incomplete read coverage will result in the assembly of discrete regions of the same transcript that are treated as independent (i.e. the 3’ and the 5’ regions of the same transcript may be assembled and represented as two seemingly unrelated transcripts). Indeed, this is one reason for the large number of transcripts in the collection. The discrete but related transcript assemblies will usually have strong sequence similarity to the same Mus gene, and will both appear in the differentially expressed gene lists. When redundant entries were encountered in parsed lists for visial display (i.e. the top 20 up regulated genes and down regulated genes – TableB1-3, Figure 6, supplemental figure 4) the highest log2FC entry was kept, while the related redundant entries were omitted.

## Acknowledgements

AP was supported by NIH R25GM115298 to Dr David Julian, TP was supported by a NSF grant 1558017 to MM and James Monaghan, ES was supported by an NSF grant 1701714.

## Supplemental Material

### Supplemental Figures

<u>Figure S1</u>. Principle Component Analysis using the prcomp function built-in to R enables visualization of sample/treatment variation in gene expression between proximal and distal skin across all significantly differentially expressed genes.

<u>Figure S2</u>. Volcano plot of log2-fold change (logFC) vs log average expression (logCPM) of gene expression data. Cut-offs designate log2FC> 2 and pvalue < = 0.05(- log10 transformed).

<u>Figure S3</u>. Heatmap of all significantly (pval>=0.5) expressed genes that exhibit a minimum log2 fold change of 2 between proximal and distal tail samples. <HEATMAP_Trinity_gene_level_DE_RESULTS_SIG.2lfc_min_zoomable.pdf>

<u>Figure S4</u>. A non-clustered heatmap of the top 20 up and 20 down-regulated genes data displaying paired proximal-distal samples side by side, and gene order from top up to top down (same order as Table 1). Thei visual representation of gene expression data further illustrates the intersample variation (proximal and distal tail skin gene expression in *Acomys* individual 1 vs. proximal and distal skin gene expression in *Acomys* individual 2), but makes clear that the trends in gene expression between distal and proximal paired samples are consistent between replicate samples.

### Supplemental Tables

<u>Table S1.</u> Raw read count data of proximal and distal *Acomys* tail skin RNA-Seq samples mapped to the *Acomys* transcriptome (GSE11308).

<featurecounts_Acomys_Distal_proximal.xlsx>

<u>Table S2</u>. Reads mapped to the Trinity based transcriptome assembly reduced from transcript ‘isoform’ level representation to ‘gene’ level representation. Thus counts associated with isoforms from the same gene have been aggregated to represent counts/gene and these data used to assess differential gene expression with EdgeR using a ‘paired’ sample design.

<Trinity_gene_level_osteo_SAMPLE_ReadCounts.xlsx>

<u>Table S3</u>. Genes identified to be differentially expressed between distal and proximal tail skin. Differential analysis was conducted with EdgeR and Benjamini & Hochberg false discovery rate correction (pval <=0.05) was applied.

<Trinity_gene_level_osteo_SAMPLES_RENAMED_AllSig-ANNOTATED.Xlsx>

<u>Table S4</u>. Significantly (pval <=0.05) differentially expressed genes that exhibit a minimum of a log2 fold-change between distal and proximal skin samples.

<Trinity_gene_level_DE_RESULTS_SIG.2lfc_min_cpm_and_FC-ANNOTATED.xlsx>

