## Supplementary figures and images for "Insights into the evolution of dermal armour: osteoderms in a mammal, the spiny mouse, *Acomys*"

### Supplemental figure 1

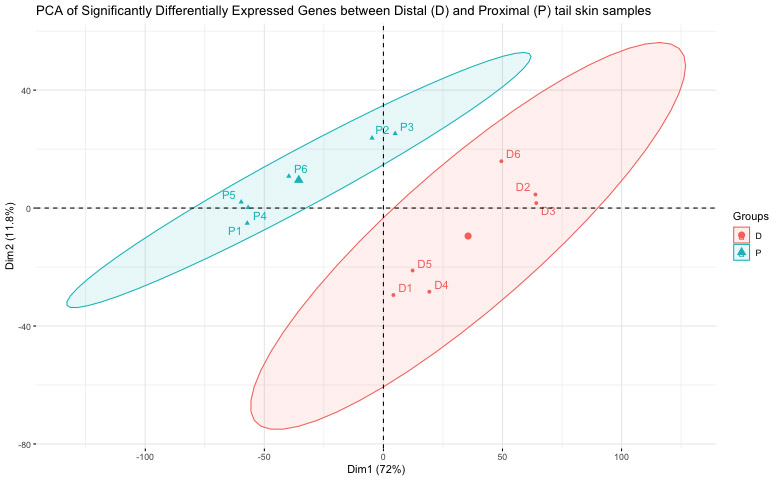

### Supplemental figure 2

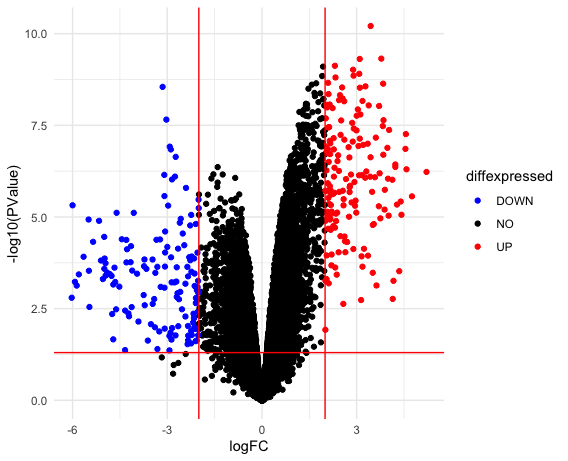

### Supplemental figure 3

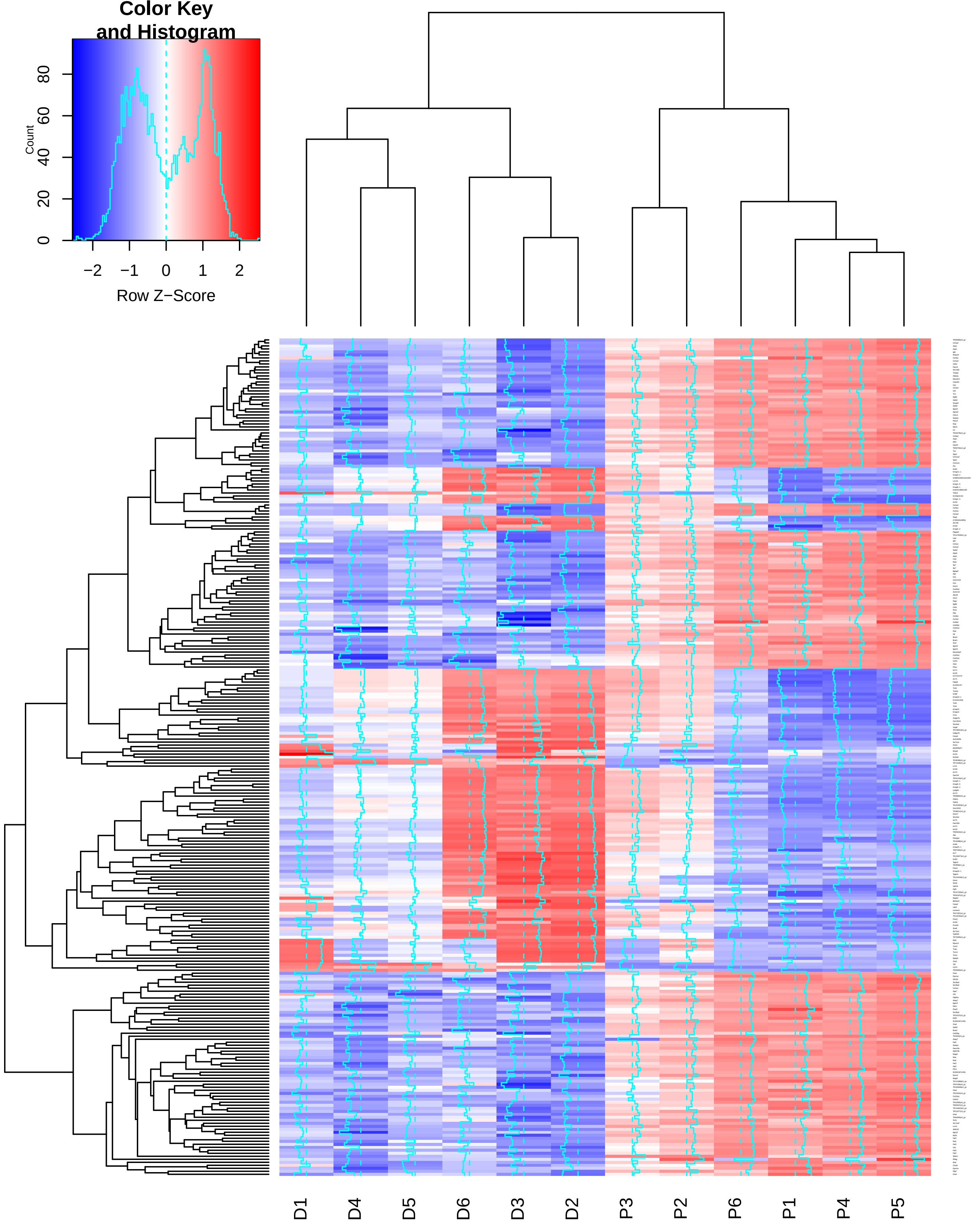

### Supplementnal figure 4

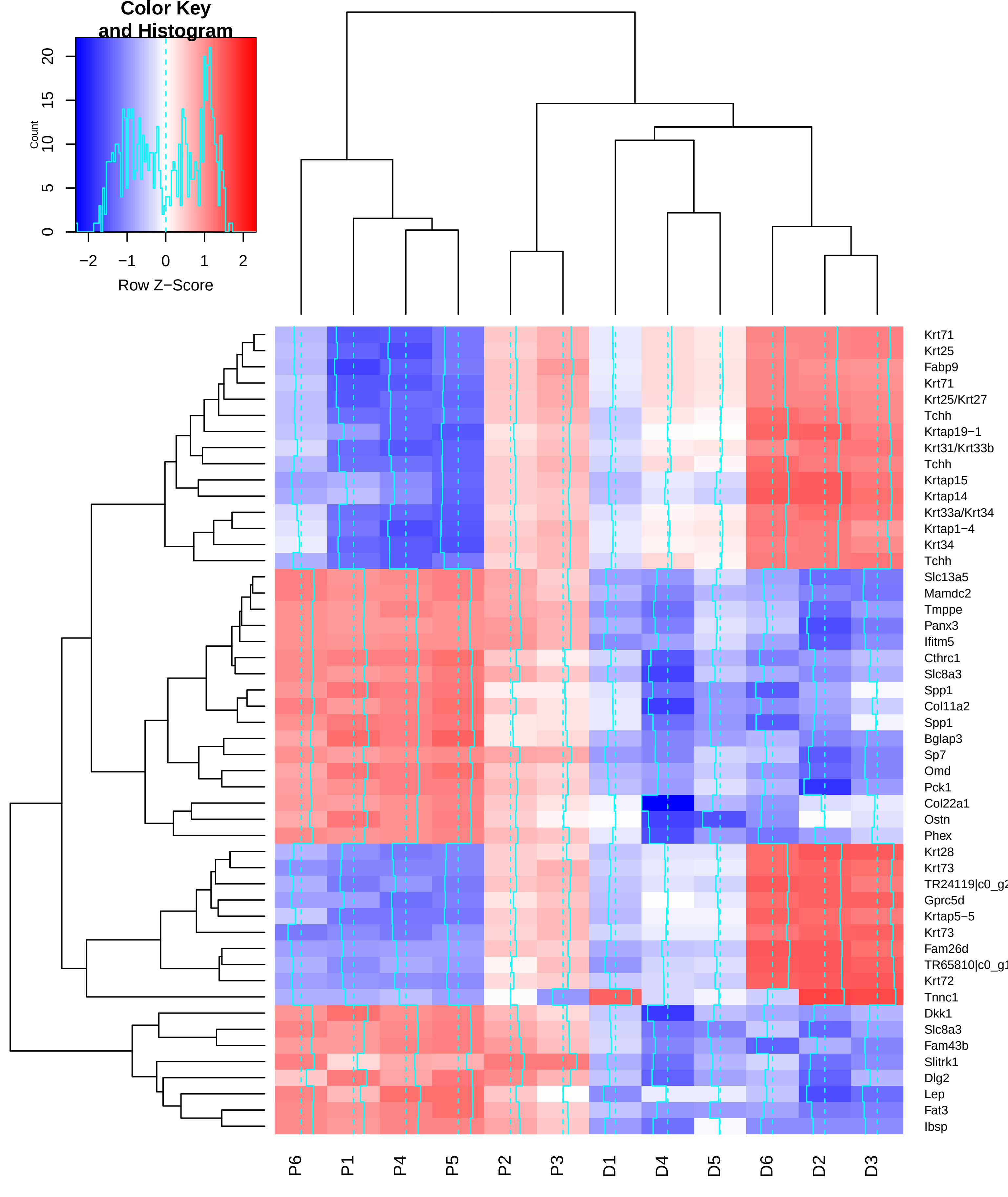
